# A comparison of Species Distribution Modeling approaches for an under-sampled parasite of public health importance, *Echinococcus multilocularis*

**DOI:** 10.1101/068734

**Authors:** Heather M. Williams, Brian Egan, Katharina Dittmar

## Abstract

**Background:** Species distribution models (SDMs) have an important role in predicting the range of emerging and understudied pathogens and parasites. Their use, however, is often limited by the lack of high-resolution unbiased occurrence records. *Echinococcus multilocularis* is a parasitic cestode of public health importance which is widely distributed throughout Eurasia and is considered an emerging threat in North America. In common with many parasite species, available data for *E. multilocularis* occurrence are spatially biased and often poorly geo-referenced.

**Results:** Here we produce three separate SDMs using MaxEnt for *E. multilocularis* using varying complexities of sampling schemes and environmental predictors, designed to make the best possible use of non-ideal occurrence data. The most realistic model utilized both derived and basic climatic predictors; an occurrence sampling scheme which relied primarily on high resolution occurrences from the literature and a bias grid to compensate for an apparently uneven research effort. All models predicted extensive regions of high suitability for *E. multilocularis* in North America, where the parasite is poorly studied and not currently under coordinated surveillance.

**Conclusions:** Through a pragmatic approach to non-ideal occurrence data we were able to produce a statistically well supported SDM for an under-studied species of public health importance. Although the final model was only trained on data from Eurasia, the global model projection encompassed all known occurrences in the United States. The approach defined here may be applicable to many other such species and could provide useful information to direct resources for future field based surveillance programs for *E. multilocularis* in North America.

## Background

Species distribution modeling is a correlative technique which pairs geo-referenced species occurrence records with maps of environmental conditions to produce a predicted distribution for the species in the wider geographic area [1]. Species distribution models (SDMs) have previously been used to model a range of human parasites and pathogens [2-5]. These models can prove useful tools for public health planning as they can help allocate resources to the areas at highest risk of infection and plan for potential changes in parasite range due to climate change or species introductions [6, 7].

The ideal occurrence data for constructing an accurate SDM spans the entire geographic extent of the species’ distribution, is free of spatial collection bias and is accurately geo-referenced at a high spatial resolution [8]. This situation is extremely rare, even for well-studied groups and almost unheard of when working with under-studied taxa such as parasites [9, 10], where data are still opportunistically collected and often geo-referenced to large political units rather than precisely co-ordinated. Perhaps paradoxically, this imperfect occurrence data makes the potential utility of SDMs even greater, to help fill in gaps in our knowledge. One such under-sampled parasite is *Echinococcus multilocularis*.

*E. multilocularis* is a widely distributed heteroxenous cestode (Cestoda, Taeniidae), which naturally exploits a sylvatic predator-prey system, with small rodents (primarily Arvicolidae and Cricetidae) as intermediate hosts, and predominantly canids (but occasionally felids) as definitive hosts [11]. Humans may become accidental intermediate hosts via ingestion of viable eggs by way of contaminated food or water, or by contact with contaminated surfaces or feces [11]. The pathology of the ensuing zoonosis is caused by the larval stage, and is increasingly recognized as a public health concern, not only in rural but also in urban and suburban areas [12]. Once established in a human host, *E. multilocularis* may cause alveolar echinococcosis (AE), which is characterized by a long-term asymptomatic onset during which time the larvae cause proliferative vesicles to form in liver tissue. If left untreated, tumors can form and infection spreads to adjacent tissues and organs [13, 14]. The advanced clinical stage of AE often corresponds to hepatic dysfunction, and the disease may be lethal, even with advanced treatment options [15]. Although 95% of an estimated 2 – 3 Million cases of human echinococcosis are caused by the closely related *E. granulosus*, *E. multilocularis* is more pathogenic, difficult to treat, and has a higher mortality rate [16].

*E. multilocularis* is typical of many widely distributed parasites in that records of its occurrence are spatially biased, patchy and often at poor resolution. Although the parasite is known to occur throughout large parts of Europe, Asia and North America [11], its cosmopolitan distribution is not truly reflected in the available records. Despite signs that the parasite is spreading in the United States and Canada [12], monitoring efforts in the Americas is uncoordinated, and have produced fewer than 50 georeferenced records (Figure S1). Furthermore, while high resolution records are relatively abundant in Western and Central Europe where sampling has been extensive, occurrences in Russia and Central Asia are often only geo-referenced at relatively large political units (Table S1). Perhaps because of this data limitation, no SDM currently exists in the literature for this species. However, with signs that the distribution of the parasite is spreading in parts of its range, the utility of a SDM which can predict the limits of that range is clear [12].

In this paper we have compiled a comprehensive database of *E. multilocularis* occurrence from literature records and used this to produce a well-supported map showing the approximate known distribution within Eurasia. We attempt to overcome the limitations inherent in the current *E. multilocularis* occurrence data with a comparison of three different SDM modeling approaches, with layers of complexity being added in each model. All three models focus primarily on predicting the distribution of the egg stage of the lifecycle. *E. multilocularis* eggs are passed in the feces of infected definitive hosts, and constitute the only developmental stage of this tapeworm outside of a warm-blooded host. This external egg stage thus represents a crucial, and relatively vulnerable step in the tapeworm lifecycle, in which egg survival is directly dependent on abiotic factors [17]. Egg survival may, therefore, play a direct role in determining the parasite^’^s geographic distribution and transmission dynamics. The wide range of rodents and canids known to host *E. multilocularis*, and the wide distribution of those groups, means that *E. multilocularis* is not likely to be limited by the distribution of its hosts. Hosts are, therefore, not a primary focus in our modeling philosophy. However, as the parasite’s transmission dynamics may be affected by host density [18] we include derived climatic variables in our final model which may affect intermediate host availability.

In our first model we define the approximate distribution of *E. multilocularis* in Eurasia, and generate a set of random occurrence points from within that distribution to produce a global SDM based on interpolated climate data. In model 2 we use the same climatic predictors as in model 1, but apply high resolution occurrence data from the literature with a bias grid to correct the relative over-representation of West and Central European records. Finally, in model 3 we use the same approach to occurrence da ta as in model 2, but add biologically relevant derived climatic and habitat predictors to the model. All three models are built using Eurasian data and are projected globally. By comparing the projections of each model we aim to: 1) test whether it is possible to produce a plausible SDM for an under-sampled species such as *E. multilocularis*; 2) test how robust the predicted distribution is to changes in modeling protocol; 3) provide an estimate of the parasite^’^s distribution in North America where it is thought to be expanding but is considered neglected from a public health perspective; and 4) quantify the niche of *E. multilocularis* eggs to determine how its range may be limited by different climatic variables.

## Methods

### Literature Records

An exhaustive search of the peer-reviewed literature was undertaken with the search term ^“^*Echinococcus multilocularis* ^”^ and/or ^“^Alveolar Echinococcosis^”^, in combination with the names of all Eurasian countries, using Web of Science. Occurrence records were collated from these papers from all developmental stages (eggs, adults) and from all host categories (intermediate hosts, final hosts, accidental hosts). Host records which may have referred to the sympatric *E. granulosus* species were excluded. Even though our aim was to model climate suitability for the egg stage, we still included location data where only the adult life stage was documented, with the assumption that viable eggs will be released into the environment by adult tapeworms. Both point data (with associated coordinate information) and area data (often listed as geo - political units) were included in our database along with the approximate spatial accuracy of the record (Table S1). The resulting dataset of 459 records, based on 120 publications was used to create a discontinuous polygon map showing the area affected by *E. multilocularis* to the highest spatial resolution allowed by the available data from Eurasia (Figure 1). Although distribution maps made for purely illustrative purposes (line drawings) have been previously published [11, 19, 20], our approach provides a data-driven map from rigorously vetted records. The same search process was employed to collect occurrence data for the Americas. This resulted in a database of 43 records, all of which originated from the United States and Canada (Figure S1). As records for North America are so scarce, these were not used during model building, but were reserved to provide independent data with which to test global model projections.

**Fig 1.**
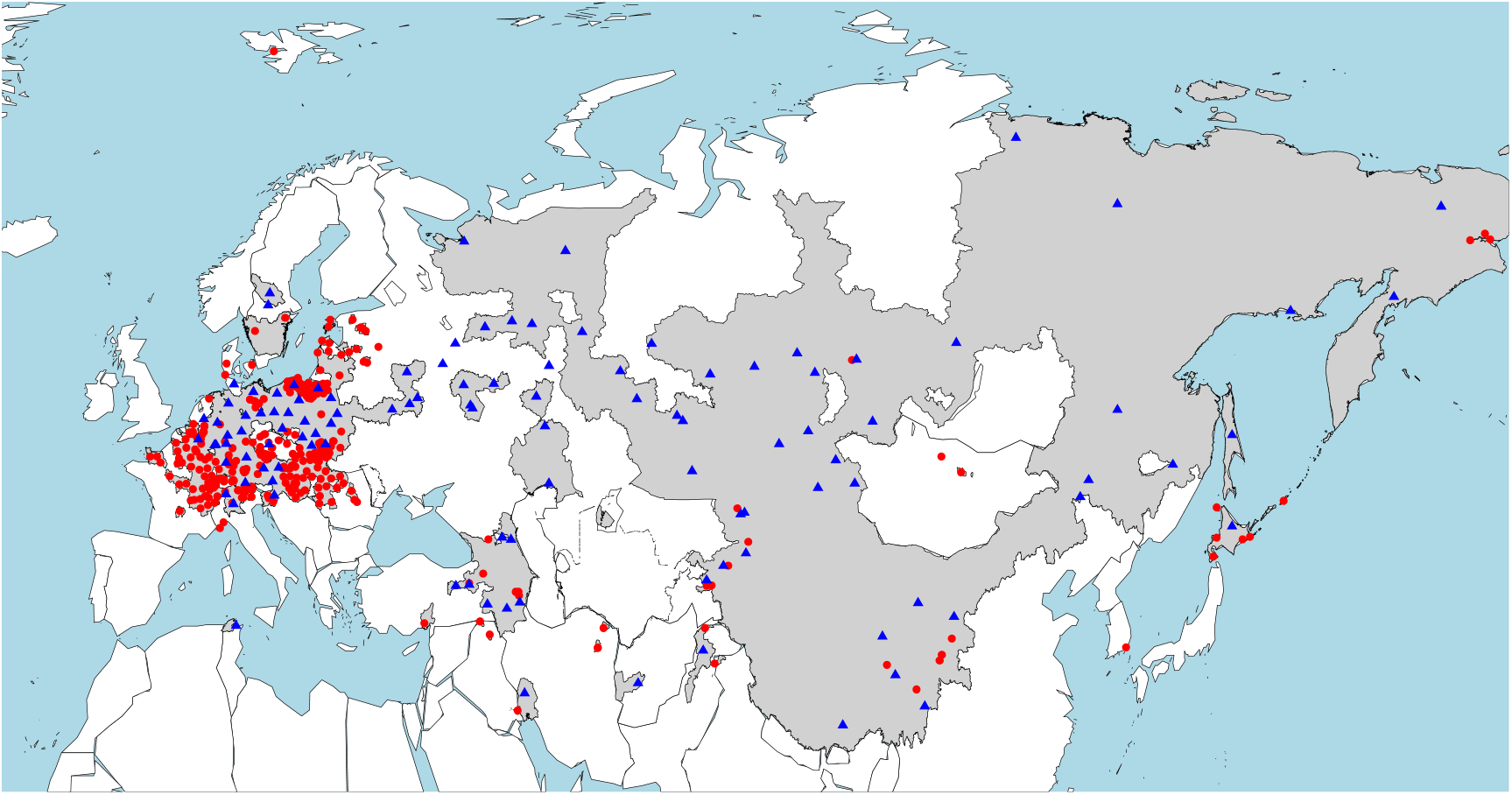
Map showing the current documented distribution of *Echinococcus multilocularis* in Eurasia. Gray regions indicate presence; red dots indicate occurrences from the literature accurate to within 10,000km^2^; blue triangles show occurrences from the literature with a resolution coarser than 10,000km^2^ (as such, their exact positioning is at random within the area they designate).

### Model 1

#### Summary

Model 1 is the most basic of the three models and uses randomly sampled occurrence data and interpolated climate data as inputs, without the use of a bias grid.

*E. multilocularis* occurrence data for model 1 was taken by randomly sampling points within the known area of Eurasian presence (i.e. the gray area in Figure 1) using ArcMap [21]. Point generation was constrained to allow only one point per 10,000 km^2^ grid cell. Five sets of 200 points were made (Figure S2) and five separate MaxEnt models were constructed to allow us to assess variability between models as a function of the exact position of the occurrence data used in the model.

Bioclim variables are one of the most commonly used data sources used in SDMs. They are publicly available globally and produced at a variety of resolutions, making them easy to use for SDM researchers. We obtained the 19 bioclim variables (Table S2) from WorldClim (http://www.worldclim.org) [22] at a 5 arc-minutes resolution and resampled to a 10,000 km^2^ grid cell size cropped to the area of known *E. multilocularis* extent using the raster package [23] in R version 3.1.2 [24].

### Model 2

#### Summary

Model 2 differs from model 1 in that the majority (165/200) of its occurrence data points are true high-resolution presences from the literature, rather than generated by random sampling. The same set of bioclimatic variables are used as in model 1, but this time a bias grid is employed.

Records of *E. multilocularis* occurrence that were geo-referenced to a resolution of at least 10,000km^2^were collected (n=165). This 10,000 km^2^ cut off point was selected as it matches the chosen resolution of our final model outputs. Where these high resolution records did not specify a coordinate, the approximate centroid of the geographic area concerned was measured and the coordinate taken. Unfortunately, very few high resolution data points are available for several large countries in which *E. multilocularis* is known to be highly endemic [11]. To compensate for this potential bias we made 5 sets of 35 random points within the area of known presence in Russia, Turkey, Mongolia and China. These 35 points were added to the 165 high resolution points to make 5 datasets of 200 occurrences, in which 35 points were different in each set. We used these five datasets to make five separate MaxEnt models to assess variability between projections attributable to the exact locations of these random points.

Even after the addition of the 35 points in Russia, Turkey, Mongolia and China, the vast majority of our occurrence points were located in Western Europe. However, as *E. multilocularis* is known to be highly endemic in Russia and parts of Eastern Europe [11], this implies a geographic bias in our dataset. To help correct this we employed a bias grid implementing the skew in research effort in the Cestoda in general. We chose to document the bias at this broader taxonomic level rather than focus on the skew in only *E. multilocularis* to help differentiate between areas of true absence and false absence. For example, we have no occurrences in our dataset from Spain, but as there is a relatively high research effort in Cestoda in Spain, we assume that our lack of records likely reflects a true absence. In Mongolia, however, we likewise have sparse records of *E. multilocularis* from the literature, but as the research effort into Mongolian cestodes is relatively low we might consider our lack of records as a false absence. To accommodate potential bias, we conducted a literature search using Thompson Reuters^’^ Web of Science with the search term ^“^cestoda ^”^ AND the name of each country within our Eurasian distribution and recorded the number of papers found (Table S3). We corrected this number by the area in km2 of each country to produce an approximation of research effort per unit area. We rasterized this data, resampled it to a 10,000 km^2^ resolution to match our environmental variables and implemented it as a bias grid in MaxEnt. As expected, research effort was much higher in Western Europe than Central Europe, Russia and most of Asia (Figure S3).

The same set of 19 bioclimatic variables as considered in model 1 were downloaded and processed.

### Model 3

#### Summary

We consider model 3 to be the most complex model. It uses the same set of occurrence data and bias grid as in model 2, but it includes derived environmental variables in addition to the 19 bioclimatic variables (model 1, 2).

As egg survival may be more directly affected by ground temperature than ambient temperature; land temperature data with a 1-degree resolution [25] was obtained with an expectation of a negative relationship between temperature and egg survival [17]. Cestode egg survival can be also limited by prolonged exposure to UV radiation [26, 27]. We thus downloaded data for monthly cloud cover at a 1 km^2^ resolution [28], with the expectation that high cloud cover may enable egg survival at higher land temperatures. Thirdly, as *E. multilocularis* eggs are known to be susceptible to drying out [17], their survival and viability may be related to both ambient humidity and the water holding capacity of soil [29]. Global data for both were downloaded at a 0.1 degree resolution [25, 30].

Where models 1 and 2 solely aimed to model regions for egg survival, additional derived environmental variables were selected for inclusion in model 3 due to a likely impact on one or more *E. multilocularis* life stages. As such, Leaf Area Index (LAI) and Net Primary Productivity (NPP) layers were included. They broadly reflect the degree of habitat vegetation and are thus expected to positively correlate with rodent abundance [31]. They were included in model 3 at a 1 degree resolution, using data provided by the MODIS Land Science Team[25]. Rodent abundance (and thus *E. multilocularis* prevalence) may also be positively correlated with habitat heterogeneity, as rodent populations are often highest in edge habitat [32]. To that effect we downloaded landcover homogeneity data at a 12.5 arc-minute resolution [33].

All layers were resampled to our model resolution of 10,000 km^2^. Summary layers were made for the variables which were available as monthly layers (LAI, NPP, Cloud cover, land temperature) reflecting annual minimum, mean and maximum values. All data preparation was carried out using the raster package [34] in R [24].

### Modeling Algorithm and Variable Selection

The spatial extent of the background training area was limited to -20^°^W and 180W^°^ and 25^°^N and 90^°^N to avoid artificial inflation of the AUC score [35]. This geographic range encompasses the entire known distribution of *E. multilocularis* in Eurasia (Figure 1).

Pairwise correlation tests were performed on all potential environmental layers for each model (Table S4). Where groups of environmental predictors were significantly correlated with a Pearson^’^s r >0.7 only one predictor was allowed into the MaxEnt model. The predictor which attained the highest AUC score as a single model variable was retained in each case (Table S4).

MaxEnt (version 3.3.3k) was used to create a SDM for *E. multilocularis* [36, 37]. MaxEnt has been shown to be among the most consistently predictive algorithms currently available for creating SDMs [38]. As MaxEnt model output can be highly sensitive to model settings [39] we used the model selection function in ENMTools version 1.4.3 [40] to find the best supported feature type, regularization value and set of predictors according to their AICc rankings. [41] (Table S5).

After tuning with AICc, the best supported models were re-run in MaxEnt with five cross validations per dataset; they were projected globally and mean projections were calculated for each model. A summary of input data for each model is provided in Table 1.

**Table 1.** Details of input data for each of the three models. Environmental predictors shown are those retained after correlation tests and AICc model selection.

|  | Occurrence Data | Bias grid? | Environmental Predictors | Features | Regularization (Beta multiplier) |
| --- | --- | --- | --- | --- | --- |
| **Model 1** | Random sampling of 1000 points within known area of Eurasian presence (Fig 1). The 1000 points were split into 5 | No | Mean diurnal temperature range (Bio2)<br>Temperature seasonality (Bio4)<br>Minimum temperature of coldest month (Bio6)<br>Mean temperature of wettest quarter (Bio8)<br>Precipitation of wettest quarter (Bio16) | Threshold | 1.5 |
|  | groupings to assess variability between groups. |  |  |  |  |
| **Model 2** | 165 occurrences throughout Eurasia specified in the literature, plus 5 sets of 35 randomly selected points in countries of known high endemism which are poorly sampled in the literature (Russia, Mongolia, Turkey and China) | Yes | Temperature annual range (Bio7)<br>Mean temperature of wettest quarter (Bio8)<br>Mean temperature of warmest quarter (Bio10)<br>Precipitation seasonality (Bio15)<br>Precipitation of wettest quarter (Bio16) | Threshold | 1.5 |
| **Model 3** | Identical occurrence points to model 2. | Yes | Mean diurnal temperature range (Bio2)<br>Temperature annual range (Bio7)<br>Mean temperature of wettest quarter (Bio8)<br>Mean temperature of warmest quarter (Bio10)<br>Precipitation seasonality (Bio15)<br>Precipitation of wettest quarter (Bio16)<br>Minimum cloud cover<br>Maximum Annual Leaf Area Index<br>Mean land temperature<br>Landcover homogeneity<br>Altitude<br>Soil water holding capacity<br>Maximum Annual Net Primary Productivity<br>Humidity | Linear and quadratic | 1.5 |

To assess variability within each model we calculated the standard deviation of each of the 5 projections within each model and assessed the difference between training and test AUC scores [41]. In the global projections, Multivariate Environmental Similarity Surfaces (MESS) were assessed to locate geographic areas in which predictors were extrapolated beyond their training range. Model predictions may not be as reliable in these areas [42].

### Model Analysis

The fit of the resulting model was assessed using the AUC statistic, which is the most commonly used threshold independent measure of fit for SDMs [39]. Response curves for each variable included in the model were created to show the statistical relationship between these environmental covariates and the probability of *E. multilocularis* occurrence.

Threshold rules are used to convert the logistic output from MaxEnt into a binary grid showing presence and absence. For our threshold rule we used a version of the 10 percentile training threshold, refined to fit the data for each model. For model 1 where all occurrence points are from random sampling, we ranked the logistic values assigned to each occurrence point and used the value achieved by the lowest 10^th^percentile as our binary cut off (0.39). For models 2 and 3 where there was a mixture of high resolution data and some random sampling of area data, we only considered the high resolution data to define our threshold. We did this to remove the risk that a low score on any of the randomly generated points could actually indicate true absence and skew the binary output. Similar to model 1, we ranked the logistic scores for each occurrence point and used the 10^th^ percentile value as our binary threshold (0.32 for model 2 and 0.28 for model 3). Maps showing the logistic output of each map were created using the rasterVis package in R [43]. Using binary versions of the three final models, we created a composite map showing areas of agreement and variability between the models.

Projecting an SDM outside of the geographic range of its training data adds some uncertainty to its output and its validity should be checked using an independent dataset [44]. To that end, we used validated occurrence data of *E. multilocularis* records in the U.S. and Canada (Figure S1) and recorded the percentage of those points encompassed in our global projections for each model.

## Results

### Model 1

While model 1 predicted an extensive area of *E. multilocularis* egg climatic suitability in Eurasia, encompassing much of Russia and North Central Asia (Figure 2), the model failed to predict occurrence in large parts of Western and Central Europe which are known to be infected with *E. multilocularis* (Figure 1). In North America, the model predicted *E. multilocularis* occurrence across a large part of the continent from Alaska to New Foundland and spreading as far south as Utah. Under model 1, the predicted North American distribution encompassed 91% of known records in the continent.

**Fig 2.**
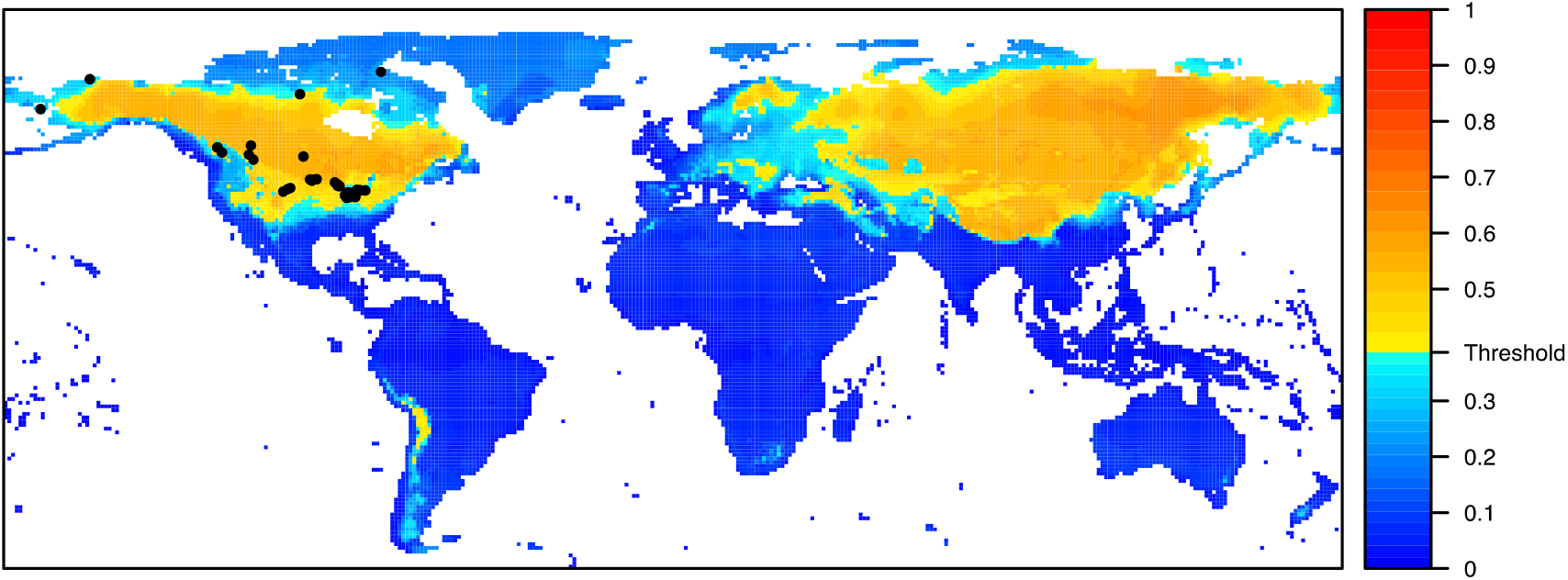
Logistic output from model 1, showing the global climate suitability for *E. multilocularis* eggs. Heatmap shows the probability of presence (between 0 and 1) with warmer colors indicating high probability of occurrence. Regions below the statistical threshold for occurrence (0.39 – see model analysis) are shown in shades of blue. Black points show known records of *E. multilocularis* in North America.

The model fit was confirmed statistically across five replicate models each with five cross-validations (mean AUC_training_ = 0.758, mean AUC _testing_= 0.697), indicating a moderately good model fit [45]. However, the drop between the training and test AUC scores could imply that the model lacks some generality and may be overfit to the training points [41].

The standard deviation between model replicates was generally relatively low (below 0.1) but was higher around the limits of the distribution including the Southern range boundary in the United States and the endemic area of Western Europe which was excluded by the averaged model (Figure S4a). However, the relatively low average standard deviation implied there was only a small difference in model output dependent on the exact location of the randomly sampled occurrence points. Analysis of MESS grids highlighted that parts of the model projection in Greenland and a long latitudinal band around the equator are outside of the model training range and may be less reliable (Figure S5a)

The projection was based on the response of *E. multilocularis* to the five statistically relevant bioclimatic variables (Table 1), three of which contributed over 80% to model fit – minimum temperature of the coldest month; mean diurnal temperature range and precipitation of the wettest quarter (Table S6). Response curves indicate the highest probability of occurrence in regions with a very low temperature in the coldest month of the year (< -20 ^°^C); with a moderate diurnal temperature range (10 – 15 ^°^C) and moderate monthly precipitation (up to 500 mm in the wettest quarter) (Figure S6).

### Model 2

The predicted distribution from model 2 covered most of Europe, with the exception of the extremes of Western and Southern Europe. It also encompassed a large latitudinal band of Northern Asia, with the exception of a part of North-western Siberia. In North America, almost all of Greenland is included within the predicted range, as well as most of Canada and the Northern United States, only excluding a narrow band along the Western coast of the continent (Figure 3). Model 2 shows a good concordance with known Eurasian records, but the predicted North American distribution encompassed just 57% of known records in the continent (Table 2) (with many of the North American points being just outside the southern boundary of predicted occurrence).

**Fig 3.**
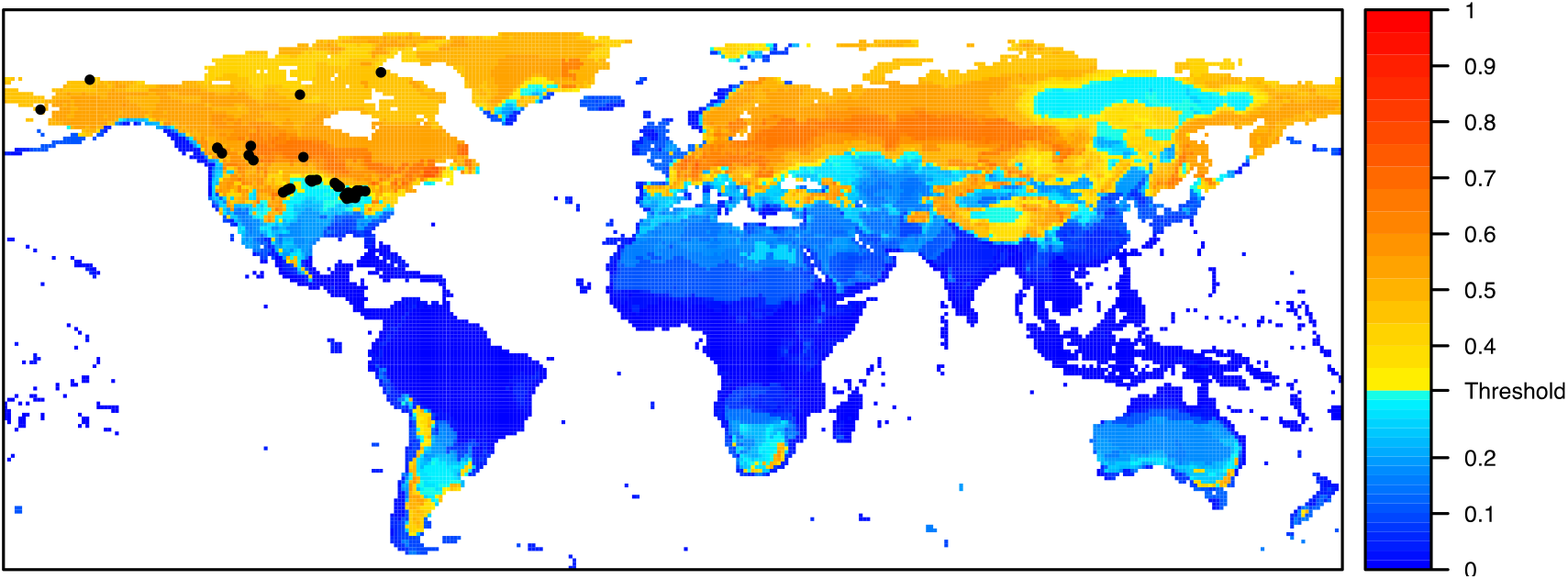
Logistic output from model 2, showing the global climate suitability for *E. multilocularis* eggs. Heatmap shows the probability of presence (between 0 and 1) with warmer colors indicating high probability of occurrence. Regions below the statistical threshold for occurrence (0.32 – see model analysis) are shown in shades of blue. Black points show known records of *E. multilocularis* in North America.

**Table 2.** Comparative metrics for the three models. AUC scores are shown as the mean value across the five cross validations of the five replicate models. Minimum and maximum AUC values are also indicated.

|  | **Model 1** | **Model 2** | **Model 3** |
| --- | --- | --- | --- |
| **Training AUC** | 0.758 (min=0.748, max=0.771) | 0.802 (min=0.7947, max=0.8078) | 0.736 (min=0.731, max=0.744) |
| **Test AUC** | 0.697 (min=0.681, max=0.711) | 0.740 (min=0.734, max=0.746) | 0.698 (min=0.689, max=0.713) |
| **Percentage of North American points encompassed in distribution** | 91 | 57 | 100 |

The model fit was confirmed statistically across five replicate models each with five cross-validations (mean AUC _training_= 0.802, mean AUC _testing_= 0.740), indicating a good model fit [45] and the highest AUC score of the three models.

The standard deviation between model replicates was generally relatively low (below 0.1) but was higher in both Greenland and the extremes of Northern Canada, and around the perimeter of the predicted area of absence in North-eastern Siberia (Figure S4b). Analysis of MESS grids highlighted that parts of the model projection in Greenland and a long, but patchy, latitudinal band around the equator are outside of the model training range and may be less reliable (Figure S5b).

The projection was based on the response of *E. multilocularis* to the five statistically relevant bioclimatic variables (Table 1), three of which contributed over 80% to model fit – mean temperature of the warmest quarter, temperature annual range and precipitation seasonality (Table S6). Response curves indicate the highest probability of occurrence in regions with a cool temperature in the warmest quarter (around 15 ^°^C); with a high temperature seasonality (> 20^°^C) and low to moderate precipitation seasonality (< 100 mm) (Figure S7).

### Model 3

The predicted distribution in Europe from model 3 was very similar to that of model 2 in that it encompassed most of Europe, excluding the extremes of Western and Southern Europe, in qualitative agreement with what is known of the distribution (Figure 4). Model 3^’^s prediction for Asia is again very similar to that of model 2, with large areas of predicted high suitability, with the exception of parts of Western Siberia. In North America, almost all of Canada and the United States are included within the predicted range, only excluding of the far West coast and Florida (Figure 4). 100% of the known *E. multilocularis* occurrences in North America fall within the predicted distribution for model 3 (Table 2). Unfortunately, due to missing data in the cloud cover dataset, model 3 is not able to make predictions for suitability for inland Greenland, parts of sub-Saharan Africa, and areas of Brazil and Bolivia (Figure 4).

**Fig 4.**
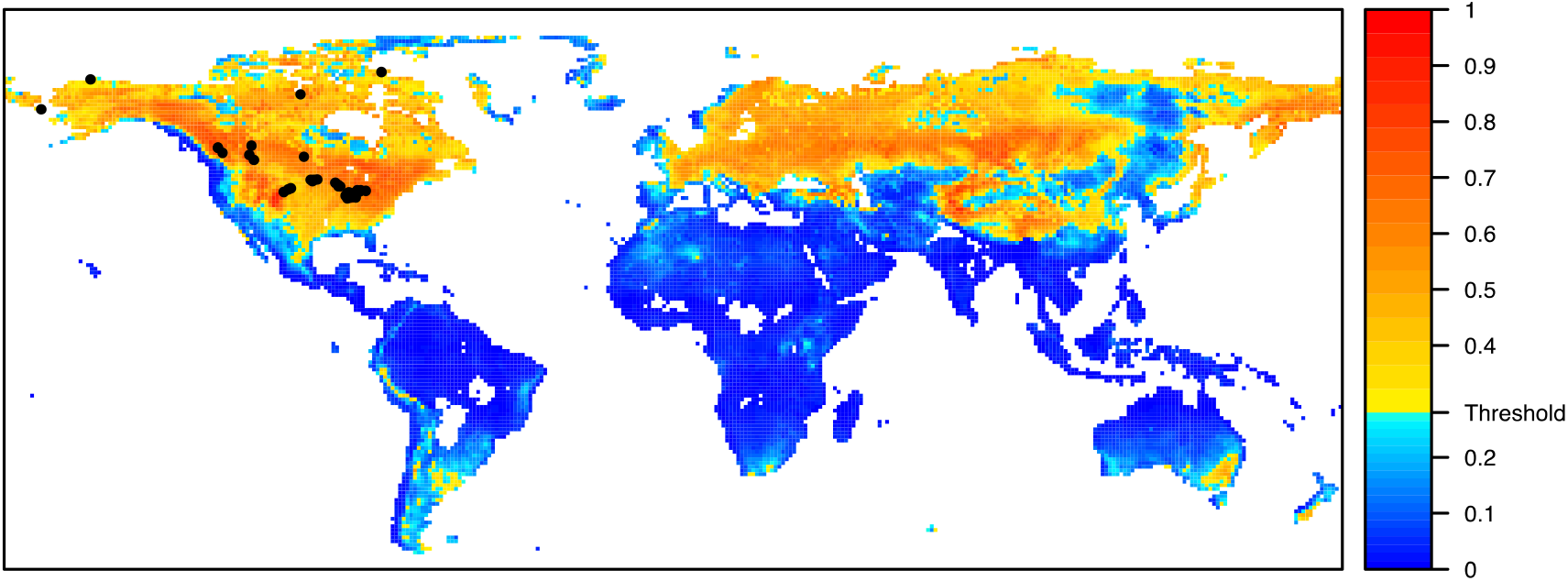
Logistic output from model 3, showing the global suitability for *E. multilocularis*. Heatmap shows the probability of presence (between 0 and 1) with warmer colors indicating high probability of occurrence. Regions below the statistical threshold for occurrence (0.28 – see model analysis) are shown in shades of blue. Black points show known records of *E. multilocularis* in North America.

The model fit was confirmed statistically across five replicate models each with five cross-validations (mean AUC _training_= 0.736, mean AUC _testing_= 0.698), indicating a moderate model fit [45].

The standard deviation between model replicates was generally relatively low (below 0.1) but was significantly higher (up to 0.3) in a geographically limited region in Western Russia (Figure S4c), meaning that there is too much uncertainty in the projection to analyze this part of the projection. Analysis of MESS grids highlighted that parts of the model projection in a long latitudinal band around the equator are outside of the model training range and may be less reliable (Figure S5c).

The projection was based on the response of *E. multilocularis* to the 15 statistically relevant bioclimatic variables (Table 1), three of which contributed over 70% to model fit – minimum annual cloud cover, landcover homogeneity and mean annual land temperature (Table S7). Response curves indicate the highest probability of occurrence in regions with moderate to high cloud cover, high landcover heterogeneity and low to moderate mean land temperature (Figure 5).

**Fig 5.**
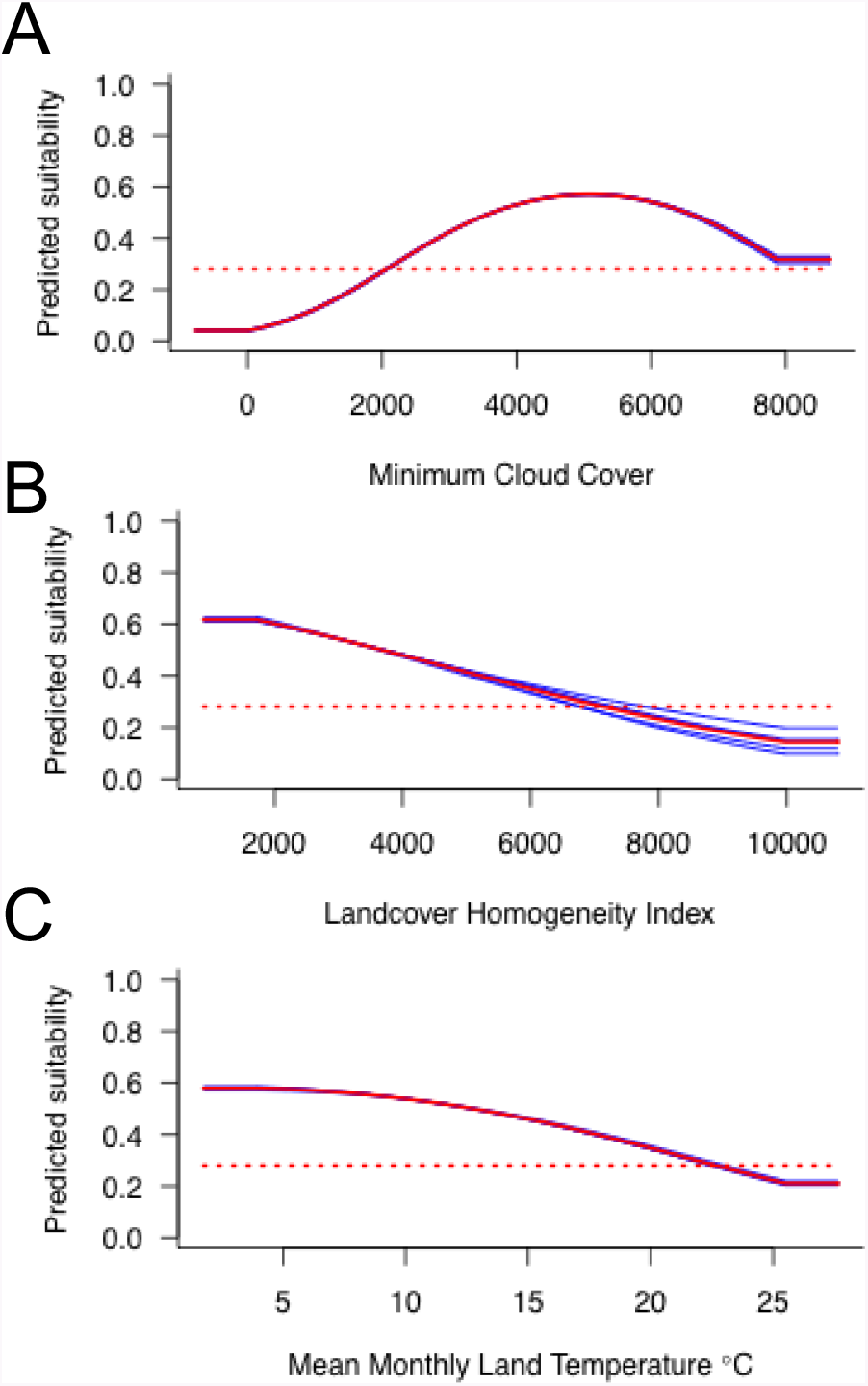
Response of *E. multilocularis* eggs to the most predictive climate variables in model 3. A. Minimum monthly cloud cover. B. Landcover homogeneity. C. Mean monthly land temperature. Blue curves show the results of each of five model cross validations, the red curve shows the average values across the cross validations. The red dashed line shows the occurrence threshold value, below which conditions are predicted unsuitable for *E. multilocularis*.

### Model Comparison

The map showing regions of agreement and conflict between the models (Figure 6) highlighted some core areas in which all models predicted *E. multilocularis* occurrence. These regions included large parts of North America, as far North as Hudson Bay, as far South as the Great Lakes in a wide latitudinal band across the continent, only excluding the West Coast. All models also predicted occurrence throughout central Russia, the extremes of North-East Russia and in parts of Northern China and Mongolia. Some parts of the northern hemisphere were not included in the projection of any of the three models. These areas include the far West of Norway, Ireland and the Western United Kingdom, Southern Europe below the Alps and the Western Coast of North America.

**Fig 6.**
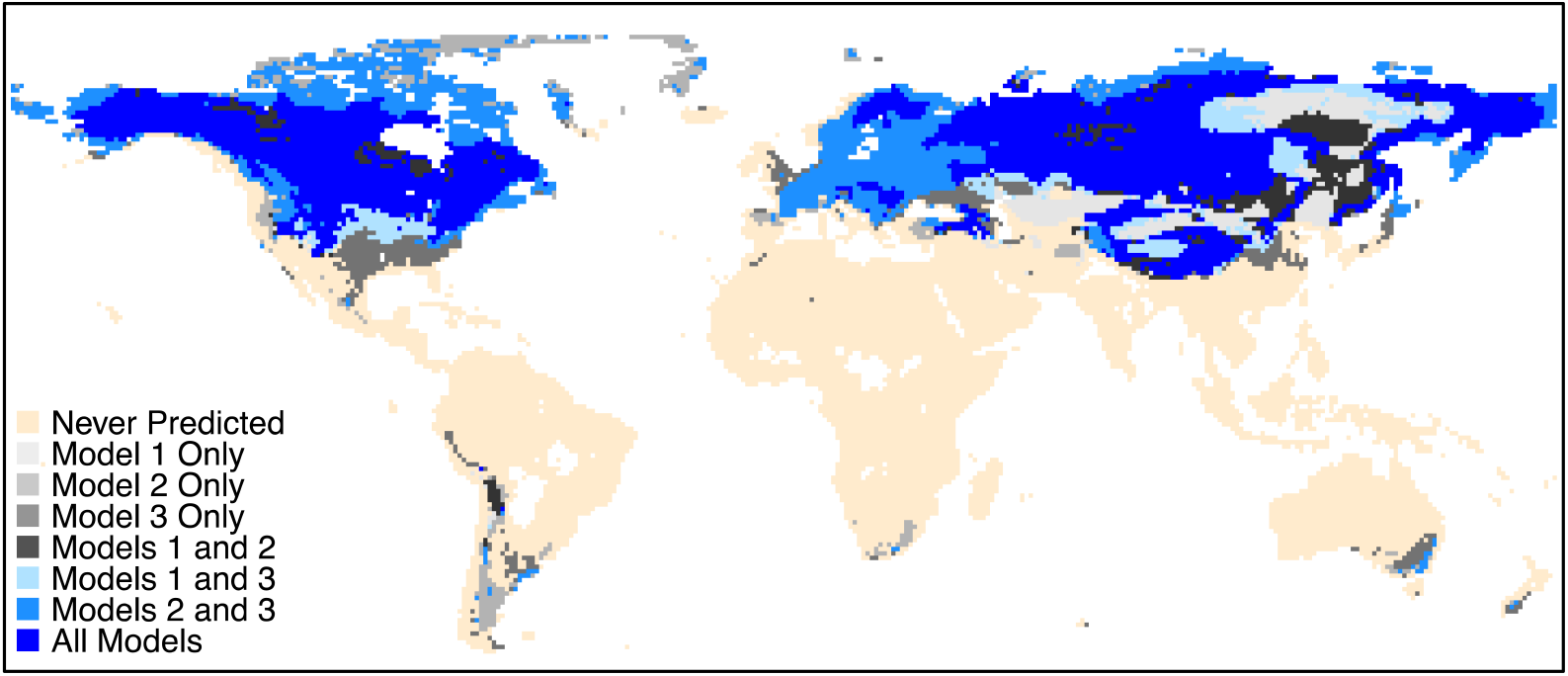
Summary map showing the areas of agreement and conflict between the three models. Regions in bright blue and predicted as suitable for *E. multilocularis* in all models. Regions in beige are never included in the predicted distribution.

The summary map shows a high degree of agreement between models 2 and 3 but poorer concordance with model 1.

## Discussion

Here we presented a test of three modeling protocols to produce a global *Echinococcus multilocularis* SDM using a challenging occurrence dataset with a strong spatial bias and patchy recording. All three models performed acceptably well according to the most frequen tly used statistical test of SDM fit, the AUC test [39]. However, while models 2 and 3 produced quite similar models, model 1 largely failed to predict *E. multilocularis* occurrence in Europe (Figure 2), despite strong empirical evidence for its occurrence. This large region of omission within the training region is most likely an artefact of an over-simplistic occurrence data sampling scheme, whereby points were randomly selected throughout the known range of the parasite (Figure 1, Figure S2). The random points not only likely introduced false positives into the dataset but appear to have driven the predicted distribution to predict only the most widespread climatic conditions encountered.

Models 2 and 3 made better use of confirmed occurrence points from the literature, compensating for their patchy spatial distribution with a bias grid. Their resulting projections are very similar at a coarse scale. Importantly, however, model 2 failed to encompass many of the known regions of North American *E. multilocularis* occurrence within its projection (Figure3), and closer comparison of the models reveals a much more extensive potential occurrence area in North America in model 3 and a more gradated predicted response (Figure 4). It is possible that model 3 produces a better fit for North America as its derived climate and habitat predictors begin to incorporate multiple life stages of *E. multilocularis* into the model (see Material and Methods). As *E. multilocularis* is capable of infecting a wide range of common and widespread host species, overlaying host distributions on top of our projection is unlikely to exclude many regions from the prediction; but using climatic predictors to infer potential peaks of host abundance, and the effect of that on *E. multilocularis*, may have served to refine model 3 compared to model 2.

The modeling protocols we have tested here are likely applicable to many other widespread species for which occurrence data is spatially biased and patchy. Our tests imply that the most realistic models come from using true presence data with a taxonomically relevant bias grid (rather than random sampling of area data) even where there is a very strong spatial bias in data collection. Furthermore, for coarse resolution models aimed at describing the maximum possible range of a species a relatively simple model of basic climate predictors is acceptable [46]. This is in line with classic macroecological theory which dictates that climate controls species distributions at the broadest scale, with habitat structural effects impacting the regional scale and biotic effects determining local scale distributions [47].

Focusing primarily on the most informative environmental variables from models two and three, we can also infer some specifics of the climatic niche of *E. multilocularis* eggs. Their temperature distribution is best characterized as relatively cool, with a decreasing probability of occurrence with warmer summer temperatures (Figure S5a); and a peak probability of occurrence at annual mean land temperatures around 0 -5^°^C (Figure 5c). This result is in general agreement with a recent study modeling human AE prevalence in China, which found peak prevalence occurred in regions of a -5^°^C annual mean temperature [48]. That egg survival may be negatively affected by warmer summer conditions is also supported by a previous laboratory analysis of *E. multilocularis* survival along a temperature gradient [49]. We also expected that egg viability may be limited by UV radiation [26]. Accordingly, we found that *E. multilocularis* appears limited by low minimum annual cloud cover (Figure 5a).

Model 3 also highlighted a relationship with seasonality, whereby *E. multilocularis* is more often found in regions with a large variation in their annual mean temperature (Figure S5b). This relationship may actually have more to do with intermediate host population cycles affecting *E. multilocularis* metacestode abundance than with egg survival *per se*. A highly seasonal environment may be prone to cyclic population explosions of rodents, leading to periods of increased *E. multilocularis* transmission when rodent population densities are high [50]. The negative relationship we found in model 3 between *E. multilocularis* occurrence probability and landcover homogeneity (Figure 5b) may also be explained by rodent population densities. Rodents are most abundant in edge habitat [32], so areas with a high landcover heterogeneity support more rodents and may increase *E. multilocularis* transmission risk. Finally, our analysis found a negative relationship between precipitation seasonality and *E. multilocularis* occurrence (Figure S5c). This apparent avoidance of a tropical/monsoon type rainfall regime may reflect a need for the eggs to both avoid desiccation in the dry season and becoming waterlogged in the wet season.

Of course, consideration of our changing climate may alter the geographic distribution of this parasite in coming years [50]. In very general terms, the *E. multilocularis* distribution may be expected to contract northwards under contemporary climate change models [50], perhaps relieving the warmest reaches of the distribution from the threat of infection. However, as relatively wide ranges of temperature and precipitation values are predicted to be above the threshold of parasite occurrence (Figures 5, S6, S7), we do not necessarily expect large range changes for this species unless the changing climate interacts with other factors affecting its distribution.

*Echinococcus multilocularis* in North America has previously been described as the ^“^great unknown^”^ due to a lack of an effective monitoring system and subsequent knowledge about its potential impact and distribution [12]. One of the objectives of this study was to help fill this knowledge gap by modeling global climate suitability for *E. multilocularis* eggs (including North America). Our models all predict the potential for parasite egg occurrence across a large latitudinal band of North America (Figure 6). This prediction not only encompasses locations where transmission has already been recorded, but also predicts well beyond those boundaries. This implies that either these regions are already infected with *E. multilocularis*, but a lack of surveys have failed to identify it; or that these areas are not currently infected with *E. multilocularis*, but would be climatically suitable for its expansion. Either scenario could pose a risk for public health.

Although there are no known records of *E. multilocularis* in the Southern Hemisphere, our model highlighted several geographically limited regions that may be climatically suitable for survival of its eggs – the Andes Mountain region of South America; the Southern tip of South Africa; South-Eastern Australia and Southern New Zealand. While New Zealand^’^s lack of a wild canid population and relatively stringent requirements for anthelmintic treatment for imported pets [51] may prevent *E. multilocularis* from ever taking hold there, South America; Australia and South Africa may be more vulnerable. The Andes region is home to not only rodent species (Cricetidae) to serve as intermediate hosts, but also possible definitive hosts (e.g culpeo - *Lycalopex culpaeus*) that could maintain a sylvatic transmission cycle. Australia and South Africa, likewise, both host the rodents and canids (e.g. Dingo (*Canis familiaris dingo*) and red fox (*Vulpes vulpes*) in Australia and black-backed jackal (*Canis mesomelas*) bat-eared fox (*Otocyon megalotis*) and Cape Fox (*Vulpes chama*) in South Africa) needed to support the *E. multilocularis* lifecycle. These areas should therefore be considered at risk, as tapeworm infections can spread quickly and jump areas of unsuitable habitat via pet and livestock trade [52].

Our model focused primarily on the climatic requirements of *E. multilocularis* eggs to produce a distribution model at the global scale. In this respect our projections could be considered a ^‘^worst case scenario’ of the potential distribution of the parasite, because limitations on the intermediate and final hosts may eventually contract this estimate. Field validation of the model projections would add certainty to these findings, and areas at the edge of our predicted distribution may yield useful data on egg tolerance limits. Future studies could also take a landscape scale approach to incorporate other factors that may limit *E. multilocularis* at a finer spatial scale. Likely limits may include land cover, proximity to water, canid population density and rodent species assemblages and density [29, 48]. Public health models may also consider how human behavior interacts with ecological factors to determine which sectors of the population are most at risk. For example, dog ownership; farming, and hunting have all been linked to increased infection risk [11, 53].

Even without these refinements, our model may already serve as a guide to show in which North American regions surveillance for *E. multilocularis* needs to be increased. It is highly recommended that suitable areas begin surveillance as soon as possible in order to provide a more accurate picture of where the parasite currently exists, as well as the directions and speed at which it could be spreading. The public in climatically suitable regions should also be made more aware of the risks posed by this disease. Information should be distributed on the risk factors associated with infection (e.g. farming, hunting, pet ownership), as well as personal preventative methods (e.g. washing hands and vegetables, avoiding contact with wild canids, deworming pets). Suburbia could be an especially vulnerable area, as an interface between wildlife and people [54], and extra efforts should be focussed in these areas.

## Conclusions

The ideal resource for building a species distribution model will always be comprehensively sampled data, free from spatial bias and accurately geo-referenced. However, given the rarity of such data sets in parasitology and disease ecology, and the high potential utility of SDMs to assist those allocating surveillance resources in these fields it may be incumbent on researchers to do the best they can with the data that is available. In this paper, a pragmatic use of non-ideal occurrence data was used to produce a statistically and empirically supported SDM for *E. multilocularis* at a coarse resolution, in which the model projections were relatively robust to minor changes in modeling framework. While data limitations may prevent high resolution SDMs for many such widespread parasites, a coarse scale climate-based model can still be a useful tool for public health planning where infection shows signs of spreading to new locations. The predicted *E. multilocularis* distribution in North America produced here may be used to direct monitoring efforts for the continent as the parasite continues to expand its range.

## Declarations

### Ethics approval and consent to participate

Not applicable

### Consent for publication

Not applicable

### Availability of data and material

All datasets used in the model are included in supporting materials with this paper.

### Authors’ contributions

HMW led the modeling process and wrote the manuscript. B E led the data collection phase of the study and assisted in the modeling process. KD was instrumental in project conception and guided the project. All authors contributed to the manuscript editing process

## Acknowledgements

We are grateful to Adam Wilson at SUNY at Buffalo (Dept. of Geography) for invaluable input on the project.

